# Metax: A Coverage-Informed Probabilistic Framework for Accurate Cross-Domain Taxon Profiling

**DOI:** 10.64898/2025.12.04.692287

**Authors:** Z.-L. Deng, N. Safaei, A. C. McHardy

**Affiliations:** Computational Biology of Infection Research, Helmholtz Centre for Infection Research, Braunschweig, Germany; Centre for Individualised Infection Medicine (CiiM), a joint initiative of the Helmholtz Centre for Infection Research (HZI) and Hannover Medical School (MHH), Hannover, Germany; Braunschweig Integrated Centre of Systems Biology (BRICS), Technische Universität Braunschweig, Braunschweig, Germany; German Center for Infection Research (DZIF), partner site Hannover-Braunschweig, Braunschweig, Germany; Cluster of Excellence RESIST (EXC 2155), Hannover Medical School, Hannover, Germany

## Abstract

Metagenomic taxonomic profiling is essential for characterizing microbial community composition in both environmental and clinical contexts. Existing profilers have greatly advanced community characterization; however, achieving accurate profiling across all domains of life - especially for archaea, fungi, and viruses - and for low-biomass, host-dominated samples, remains challenging. We describe Metax, a cross-domain taxonomic profiler that employs probabilistic modeling of genome coverage to distinguish true community members from artifactual signals arising from reference contamination, local genomic similarity, or reagent-derived DNA fragments. In comprehensive benchmarks across more than 500 samples, Metax demonstrated accurate species-level profiling, with consistent performance for bacteria, archaea, eukaryotes and viruses, and robustness to shallow sequencing. Applied to an oral microbiome cohort, Metax identified differentially abundant viral taxa distinguishing peri-implantitis from healthy sites, while analyses of tumor microbiome data revealed reagent-borne contaminants and potential reference misassemblies. By integrating coverage-informed statistics, Metax delivers accurate, robust, and interpretable cross-domain taxonomic profiles, maintaining stable performance across diverse sequencing depths and sample types.

## Introduction

Shotgun metagenomics enables the direct study of microbial communities through sequencing of environmental or host-associated DNA^1–4^. This approach allows the simultaneous detection of bacteria, archaea, viruses, and microeukaryotes without requiring cultivation, and has become indispensable for characterizing community composition^5–7^, function^8–10^, ecological interaction, and host–microbe relationship across environments^11^.

A wide range of analytical techniques has been developed to interpret metagenomic data. Assembly reconstructs contigs from mixed-community reads, which are subsequently grouped through binning into metagenome-assembled genomes (MAGs). These genome-resolved approaches provide valuable genomic context for taxonomical, functional, and evolutionary analyses^1,12^. Taxonomic profiling methods allow to determine the taxonomic identities and relative abundances of community members at different ranks. The comprehensive quantification of taxa abundance by profiling approaches is essential for large-scale ecological and clinical studies^13–15^. Despite major progress over the past years, challenges remain^16,17^. Ambiguity in read assignments^18^, insufficient genome coverage due to shallow sequencing or limited availability of microbial biomass^19,20^, and reference misassemblies^21^ compromise species-level resolution and abundance estimation. These issues are further exacerbated in low-biomass or host-dominated samples, where reagent-derived DNA (“kitome”) and residual host sequences can obscure genuine microbial signals^22,23^.

To address these challenges, we developed Metax, a cross-domain taxonomic profiler for accurate detection and abundance estimation of taxa from shotgun metagenomes using genome coverage information within a probabilistic framework. We systematically evaluated Metax on a range of datasets representing common application scenarios, including deeply sequenced complex environments and shallow sequence data from clinical specimens covering multiple body sites, and cohort studies, as well as long and short sequence data. To demonstrate further applications of Metax, we applied its’ coverage-based statistics to identify likely artifacts in genome assemblies and distinctive, interpretable taxon signatures of periodontitis covering multiple taxonomic domains.

## Results

### The Metax approach for taxonomic profiling using coverage information

The key concept underlying Metax is to combine sequence homology signals with genome coverage metrics—specifically the breadth and depth of coverage— in a statistical framework to achieve accurate taxonomic profiling. By modeling the breadth and depth of coverage, Metax quantifies the likelihood of true taxon presence, while an Expectation–Maximization (EM) algorithm refines abundance estimation by resolving multi-mapped reads. This coverage-informed probabilistic design enables Metax to distinguish genuine community members from artefactual signals caused by local genomic similarity, reference contamination, or sequencing artefacts. This coverage-informed framework filters out artifactual taxa—caused by reference contamination, kitome-derived DNA fragments, or shared conserved orthologs and mobile genetic elements—by detecting uneven read mapping across reference genomes (Fig. 1a). An EM algorithm further refines the abundance predictions. The workflow comprises five main steps listed below (Fig. 1a).

**Fig. 1.**
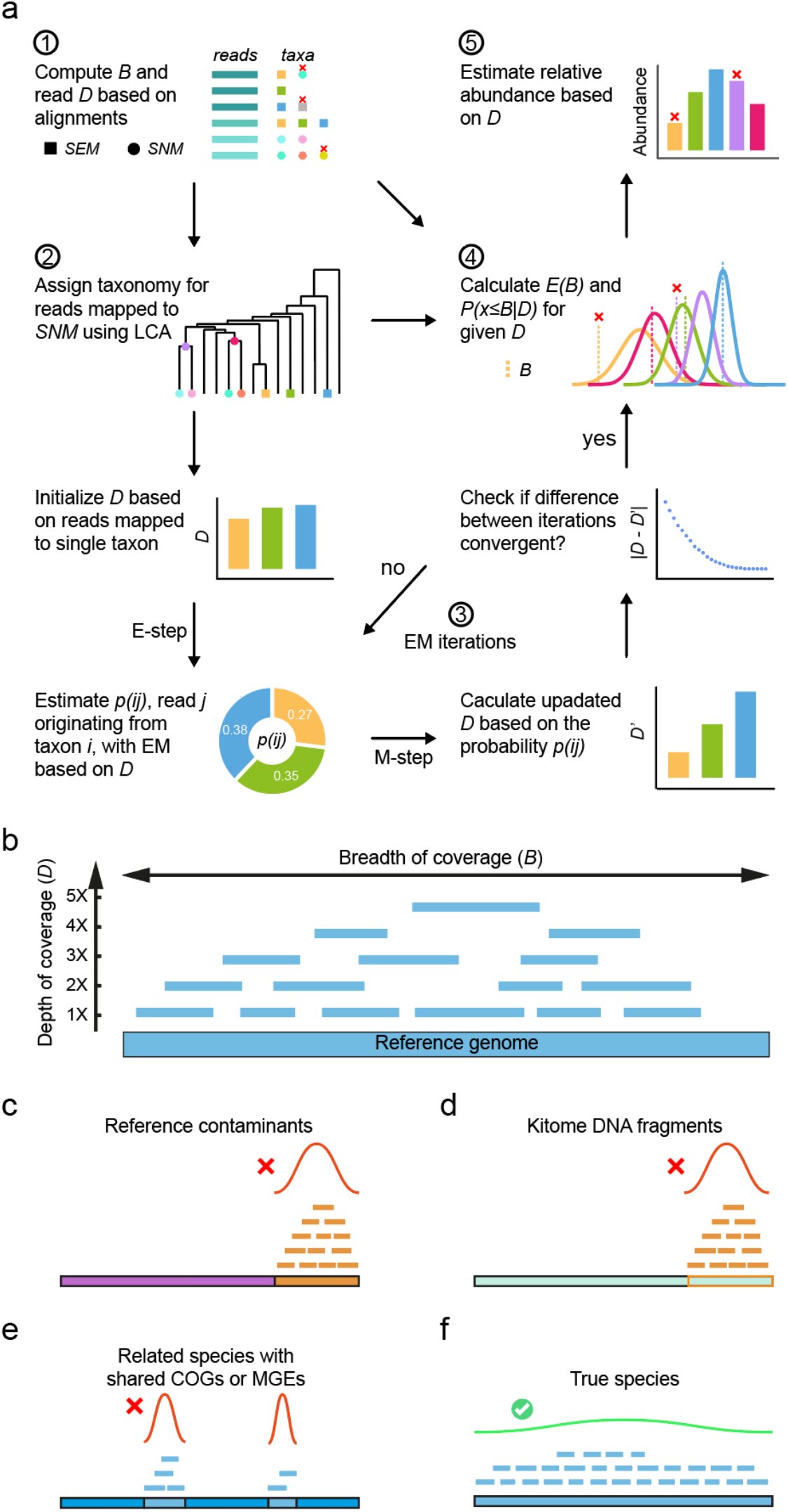
Metax using coverage information for taxonomy profiling. **a)** The Metax workflow. At step 1 and 2, the dot represents species with exclusively mapped reads (*SEM*), while square indicates species without exclusively mapped reads (*SNM*). At step 3, the *p(ij)* denotes the probability of read j originating from taxon i in the EM iteration. *B* is breadth of coverage, while *D* is depth of coverage. At step 4, *E(B)* represents the expected *B*, and the density plot is the distribution of *B* given the *D*. The observed *B* is the dashed line in each probability density plot, while *P(x*≤*B*|*D)* represents the probability of an observed *B* or smaller given the *D*. At the last step, the abundances of higher rank taxa determined by LCA are also calculated. **b)** Illustration for breadth and depth of genome coverage. Metax flags false discoveries resulting from reference contamination **c)**, kitome DNA fragments introduced in library preparation **d)**, related species with shared conserved ortholog genes (COGs) or mobile genetic elements (MGEs) to the actual species present in the data **e)**, and identifies the true taxon in the data **f)**. For b) to f), the long and wide line at the bottom shows the reference genome, while the short and thin lines are the reads map to it. In c), the orange segment in the reference genome is foreign DNA contamination included during assembly or binning. In d), the region outlined in orange corresponds to kitome-derived DNA fragments introduced during library preparation, which are reflected as contaminant reads in the sequencing data. In e), the light-blue segments denote conserved orthologous genes or mobile genetic element sequences shared between the artifactual species and the true species. In c) to f), the bell-shaped curves (red for FPs, green for TP) show the density distribution of read coverage across the reference genome, with a red cross marking the flagged artifactual signal and green checkmark indicating a true positive.

### Step 1: Genome Coverage Calculation

Metax first aligns sequencing reads to a reference genome database using Modular Aligner (MA)^24^. Alignments are filtered based on sequence identity and mapped length. After filtering, Metax calculates the observed Breadth of coverage (*B*), which represents the fraction of the genome covered by mapped reads (Fig. 1b), and Breadth of chunk coverage (*Bc*) the fraction of genome chunks covered by reads, reflecting read distribution uniformity, especially relevant in low-abundance taxa and/or shallow sequencing data. To compute *Bc*, the reference genome is partitioned into a fixed number of chunks of equal length (Material and Methods).

Short sequence reads and the inherent complexity of microbial communities often lead to multi-mapping, where a single read aligns to genomes of multiple taxa. Metax categorizes species into: species with exclusively mapped reads (*SEM*), which are species with at least one read mapping to its genome unambiguously, species without exclusively mapped reads (*SNM*), representing species for which every read that maps to its genome also maps to one or more other species’ genomes. Similarly, mapped reads are grouped into reads mapped exclusively to one species, reads mapped to *SEM* species, and reads mapped to *SNM* species.

Then, Metax computes an initial depth of coverage (*D*) from exclusively mapped reads and records read-taxon associations for subsequent probabilistic assignment.

### Step 2: Probabilistic taxonomic assignment

Metax addresses multi-mapping complexities by combining EM and least common ancestor (LCA) approaches. The taxonomy of exclusively mapped reads are directly assigned. Read mapped to several *SEMs* undergoes probabilistic assignment via EM iterations. The taxonomy of reads mapped to *SNMs* are assigned with the LCA approach.

### Step 3: EM based depth of coverage estimation

Metax employs an EM algorithm to estimate each species’ true depth of coverage. For species *i* ∈ {1, …,*N*}, let *D*_*i*_ denote the depth of coverage for species *i*. Prior to the EM optimization, Metax initializes *D*_*i*_ using only reads that map uniquely to species *i*:

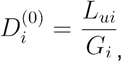

where *L*_*ui*_ is the total bases of uniquely mapped reads for species *i*, and *G*_*i*_ is the length of its reference genome.

The EM algorithm begins with the “expectation” step, where each species of origin for a multi-mapped read is assigned a probability. This is calculated by dividing the depth of this species by the sum of the depths of all species to which the read maps. Let *N* be the number of candidate species, *R* the number of reads, *D*_*i*_ the current estimated depth of species *i, M*_*ij*_ =1 if read *j* maps to species *i* and 0 otherwise, and *l*_*j*_ the length of read *j*. For each read j that maps to multiple species, the probability that it originated from species i is computed as:

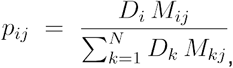

This assigns each read fractionally to the species in proportion to their current depth estimates.

The expected depth contribution of read *j* to species *i* is then:

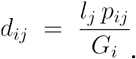

In the maximization step, species-specific depth estimates are updated by summing contributions from all reads:

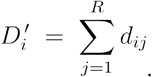

The EM updates are repeated until depth estimates stabilize. Convergence is reached when

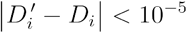

for all species *i*, indicating that additional iterations do not meaningfully change the solution.

### Step 4: Expected *B* and likelihood of observed *B* given depth *D*

The sequencing process can be modelled as random sampling^25,26^ of fragments from genomes present in a microbial community. Given a depth of coverage *D*, the expected breadth of genome coverage, *E*(*B*), is

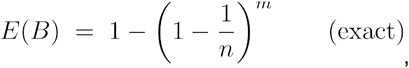

where *n* is the number of non-overlapping genome windows of size equal to the average read length, *m* is the number of mapped reads, and *D*=*m/n* is the genome depth.

Using the approximation (1 − *x*/*m*)^*m*^ ≈ *e*^−*x*^ for large *m*, this becomes

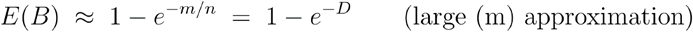

When the observed breadth *B* (computed as the fraction of genome windows covered by at least one read) falls substantially below the expected value *E*(*B*), it indicates that the reference genome does not fully represent the true underlying sequence. Such discrepancies commonly arise from contaminated or misassembled reference genomes, kitome-derived DNA fragments, or local sequence similarity that does not reflect true taxonomic presence (Fig. 1c-f).

We could also calculate the probability of the reads distribution on the genome. Given *m* mapped reads, let *X* denote the number of windows that receive at least one read. The probability of observing exactly *k* covered windows is

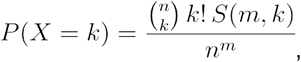

where *S*(*m,k*) are the Stirling numbers of the second kind^27^, representing the number of ways to partition m labeled reads into k nonempty windows. The Stirling term is computed as:

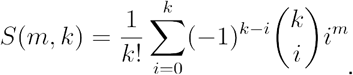

The exact (one-sided) cumulative probability of observing at most *k* covered windows is:

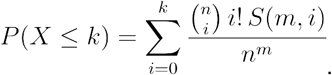

Because this calculation becomes computationally expensive for large *m*, Metax evaluates the exact probability only when *m* <100.

When the number of genome windows *n* and mapped reads *m* are large, the exact calculation becomes computationally prohibitive. However, in this situation, coverage across windows is well approximated by independent sampling events, allowing the number of covered windows to be modelled using a binomial distribution with coverage probability:

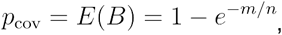

yielding the approximation

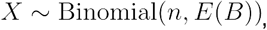

where *X* is the number of covered windows. Thus, the probability of observing a breadth as low as the observed count k can be approximated as

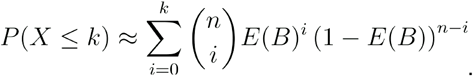

For computational efficiency, Metax uses the normal approximation to the binomial when *n* is large. With observed breadth *B* = *k*/*n*, the one-sided *p-value* for obtaining a breadth less than or equal to *B* is

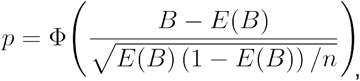

where Φ (·) denotes the standard normal cumulative distribution function.

### Step 5 Relative abundance calculation

Finally, the relative abundance for each species is computed by dividing its final depth estimate by the summed depth of coverage for all species not flagged as false positives in the previous step:

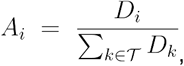

Where *A*_*i*_ and *D*_*i*_ are the relative abundance and final depth of coverage for species *i*.

### Metax improves cross domain taxonomic profiling at species rank for complex communities from human gut

Taxonomic profilers demonstrated substantial performance improvements in recent benchmarks^16^ encompassing both taxon identification and abundance estimation. Challenges remain in low ranking taxon assignment, such as at the species level, and for non-bacterial taxa, such as archaea and viruses. Due to its coverage-informed statistics, Metax is well suited for taxonomic assessment at lower taxonomic ranks. We evaluated Metax’s performance in comparison to leading profilers relying on alternative techniques such as marker-genes (mOTUs3^28^, MetaPhlAn4^15^) and k-mer sketching (Sylph^29^), and k-mer matching (Phanta^30^, Centrifuge^31^), using ten simulated human gut metagenomes. In this and following evaluations on ascites and cerebrospinal fluid (CSF) samples, Metax was run with a reference sequence collection released in 2022 (Material and Methods), to ensure consistent novelty of benchmark data relative to genomes in reference sequence collections relative to other profilers. For a straightforward comparison to recent versions of MetaPhlAn4 (version 4.1.1) and mOTUs3 (version 3.1) on the gut metagenomic dataset, we also ran Metax with a more recent reference genome collection from July 2025 (Metax_newdb) (Material and Methods).

The quality of predicted taxon profiles was assessed using the completeness and purity of identified taxa, relative to the underlying ground truth profiles^16^. Taxon abundance estimates were evaluated using the Bray–Curtis distance, and abundance rank error at species level and the weighted UniFrac error across ranks^16^—which capture deviations in abundance magnitude, taxonomically informed abundance differences, and rank-order accuracy, respectively. The abundance rank error is a metric introduced in this study, measuring how accurately a taxonomic profiler reconstructs the abundance order for a microbial community, by assessing the discrepancy between the predicted and true relative abundance rankings of taxa for a microbial community.

Gut metagenome data were simulated based on real gut taxon abundance profiles^32^ and augmented with representative fungal and viral species, including bacteriophages hosted by resident bacteria, to create realistic multi-domain communities (Material and Methods). These represent a complex microbial community sequenced with sufficient depth for reference based genome recovery (2G bp sequencing reads)^22^.

In taxon identification, both Metax and Metax_newdb improved at species rank by 4-102%, and 21-134% in F1-score compared to other profilers (Fig. 2a). This primarily stems from its high completeness of taxon recovery (0.75 ± 0.018), and ability to accurately identify viral and fungal taxa without compromising purity (0.82 ± 0.26). In relative abundance estimates, Metax performed well across all ranks (weighted UniFrac error (0.21 ± 0.047, Fig. 2a), representing a 10–63% improvement over other profilers. At the species level, Metax achieved the closest relative abundance estimates, reflected by the lowest Bray–Curtis distance (0.39 ± 0.066; 5– 47% reduction) and the lowest abundance rank error (0.21 ± 0.019; 16–73% decrease), demonstrating a notable improvement across relevant metrics for metagenomes of complex, deeply sequenced, and taxonomically diverse communities. In terms of runtime, Metax demonstrated an efficiency comparable to marker gene-based methods, by completing profiling of each 2 Gb sample in just 8 minutes, using a peak memory of 55 GB (Supplementary Fig. 1).

**Fig. 2.**
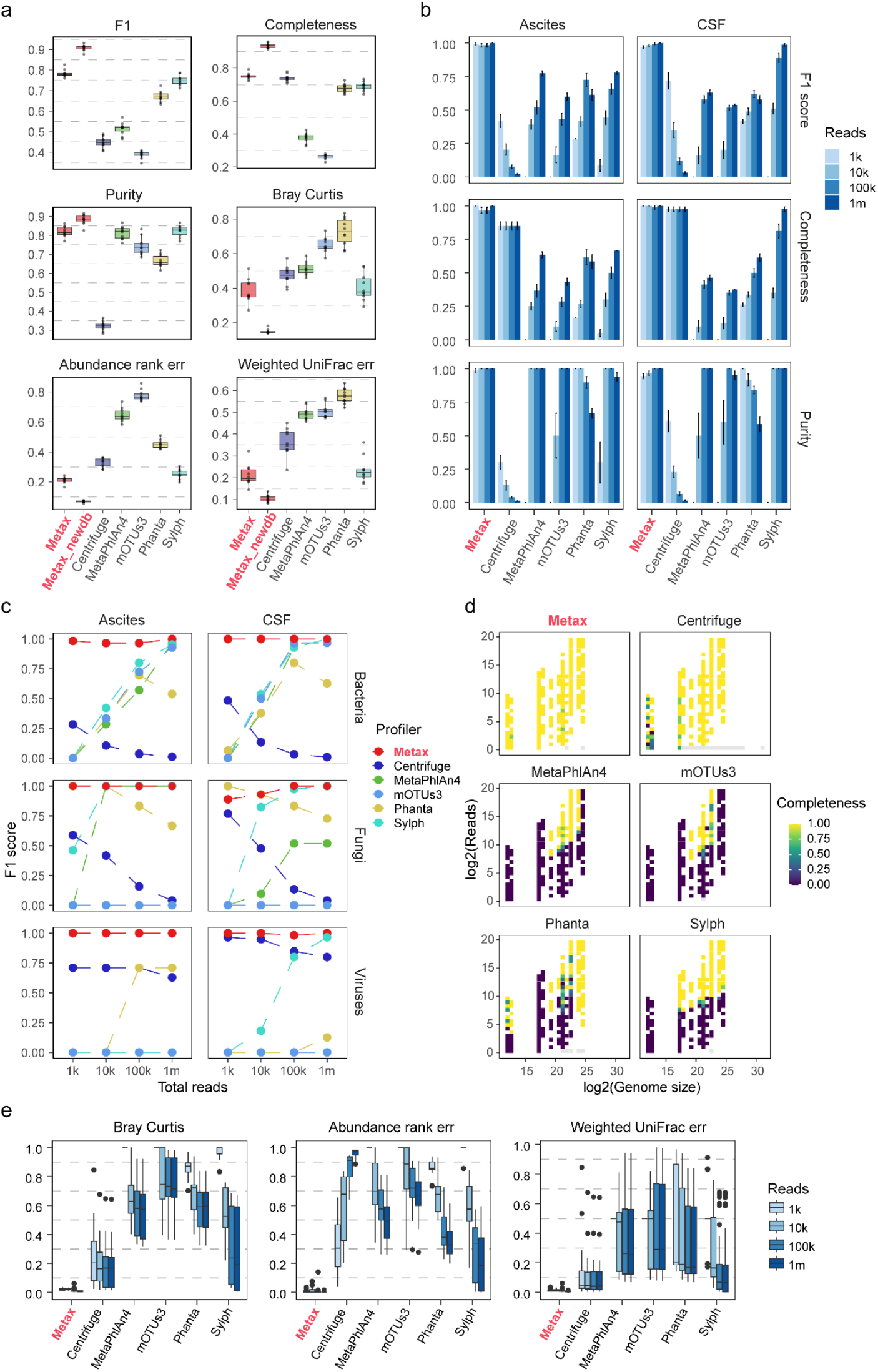
Performance evaluation of Metax on simulated gut and clinical metagenome samples at species level. **a)** Performance metrics of different profilers on gut metagenomes with around 10 million reads per sample. F1-score reflecting predicted taxon profile quality (larger is better), Bray Curtis distance and weighted UniFrac error, reflecting accuracies of abundance estimates at species level and across ranks, respectively (smaller is better), and the ‘abundance rank err’, reflecting adherence of taxon ordering (smaller is better). **b)** Quality of predicted taxon profiles for different profilers on ascites and CSF samples at lower sequencing depths (1k, 10k, 100k, 1million). **c)** F1-score of different profilers for detecting bacteria, fungi and viral taxa, respectively from ascites and CSF samples with different sequencing depth. **d)** Completeness of taxon profile recovered with different methods from the samples. The genome sizes and read counts of taxa are grouped into 30 bins shown in x- and y-axis, respectively. **e)** Abundance estimate at different sequencing depths.

### Improved performance at shallow sequencing depth

In host-associated metagenomics, for example when sequencing samples with suspected infections to support diagnostics, microbial biomass is often low relative to the host background, resulting in a limited number of microbial reads, even with host depletion^33,34^. Because host genomes are orders of magnitude larger than microbial genomes and consequently generate far more DNA fragments, low-biomass samples dominated by host DNA present another obstacle to accurate taxonomic profiling. To assess how Metax performs for varying microbial read depths and across different taxonomic groups, we simulated metagenomes from 10 infected ascites and 10 CSF clinical samples. The microbial taxa included in the simulations were selected based on those commonly reported in ascites^33,35–38^ and CSF samples in the literature^39^, respectively. Each simulated ascites sample contained 3 bacterial, 1 fungal, and 2 viral species, while each CSF sample comprised 3 bacterial, 2 fungal, and 2 viral species. Each community was then simulated for samples at four sequencing depths: 1,000 (1k), 10,000 (10k), 100,000 (100k), and 1 million (1m) microbial reads.

In taxon detection, Metax demonstrated robust performance across all sequencing depths in F1-score (0.99 ± 0.027), completeness (0.99 ± 0.039) and purity (0.99 ± 0.037) (Fig. 2b) across ascites and CSF samples, improving 1.7-22x on other methods for 1k, 2.1-5.4x for 10k, 1.3-10x for 100k and 1.1-37x for 1 million reads, and identifying all bacterial, fungal and viral taxa (Fig. 2c). The improved taxon identification by Metax is most evident for taxa with small genome sizes when using shallow sequencing (Fig. 2d). Other methods demonstrated close performances for bacterial taxa only when reaching 100k or 1 million reads, as well as varying performances for fungal taxa, which have the largest genomes, and for viral taxa. Some viral taxa were missed altogether across data sets and sequencing depths (Fig. 2c).

For abundance and abundance-rank estimation, Metax showed consistently low error at the species level, with a Bray–Curtis distance of 0.016 (13–50× lower than other profilers) and an abundance rank error of 0.0079 (65–102× lower). Across taxonomic ranks, weighted UniFrac error remained low at 0.0107, representing a 9–42× reduction relative to other methods across sequencing depths and sample types (Fig. 2e). These gains were most pronounced under shallow sequencing: at 1k reads per sample. Together, these results demonstrate that Metax delivers robust and accurate taxon profiles for samples with shallow sequencing depth, in terms of identification of the taxa present, as well as in estimating their abundances, making it well suited for such challenging applications.

### Accurate species-level taxon identification and abundance estimate for complex, deeply sequenced marine communities

To assess performance for further environments, we also evaluated Metax on the CAMI II marine short read and long read data, which correspond to complex microbial communities sequenced at substantial depth, with 5 Gb of long and short read data each per sample. For comparison with other techniques, in addition to the taxonomic profiles of tools^6,40^, Centrifuge^31^, Metalign^41^, MetaPhlAn2^42^ and mOTUs2^7^ provided in the CAMI II challenge, we ran again recent marker-gene based^28,15^, k-mer sketching^29^ and k-mer matching^43^ methods. To ensure a fair comparison, the same database version provided then was used for Metax and other methods, where applicable. Notably, for the recent marker gene based profilers their latest custom marker gene databases were used, which include more genome data underlying the CAMI II marine challenge data set than the versions used by other tools, which likely beneficially impacts performances.

In taxon identification, Metax achieved the highest F1-score (0.88) at species level, both for long and short read data (Fig. 3a), with other methods ranging from 0.34 to 0.88. In abundance estimation across ranks, Metax had a low weighted UniFrac error (0.015), similar to methods using more recent reference sequence collections, improving by 10-69% on others. Abundance estimation was particularly accurate at species level (Bray-Curtis distance 0.01; improving 16-67% relative to others). Comparison of Metax predicted abundances to the ground truth at species level yielded a Pearson’s correlation coefficient of *r* = 0.97 (two-sided p < 2.2 × 10^−16^), indicating a highly significant concordance between predicted and actual abundance profiles (Fig. 3b). It also demonstrated the best species-level abundance ranking, as reflected by the lowest abundance rank error (0.12; 11-71% improvement). Together, these results demonstrate the capability of Metax to generate accurate taxon profiles for complex communities from deeply sequenced metagenomes - with the most pronounced benefits evident for low-ranking taxa such as species.

**Fig. 3.**
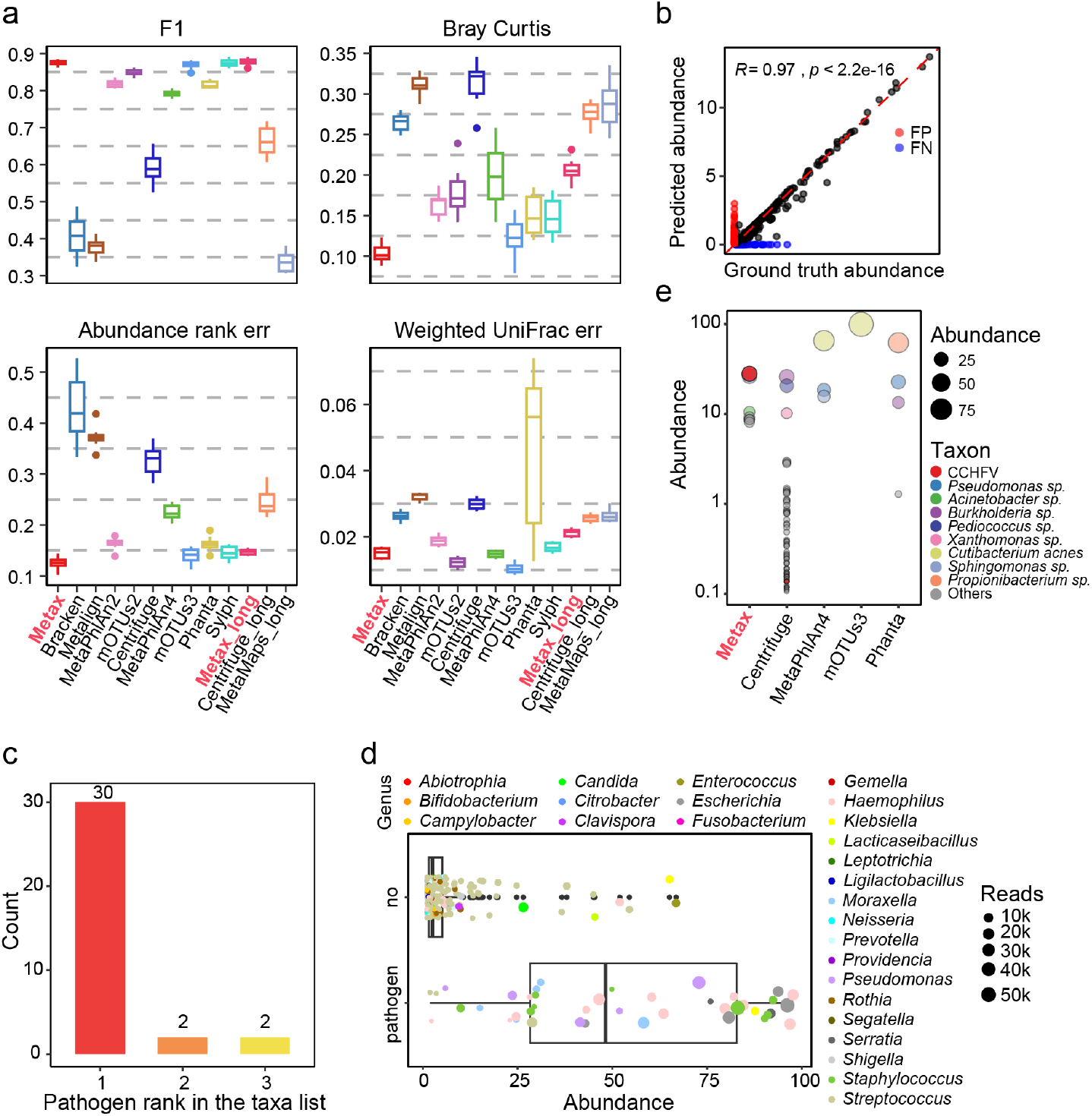
Comprehensive performance comparison of state of the art profilers on the CAMI II marine and pathogen detection datasets at the species level. **a)** F1-score reflecting predicted taxon profile quality (larger is better), Bray Curtis distance and weighted UniFrac error, reflecting accuracies of abundance estimates at species level and across ranks, respectively (smaller is better), and the Diversity diff, reflecting accuracy of diversity estimates (smaller is better) **b)** Metax taxon abundance predictions and ground truth abundances. The x-axis shows the ground truth abundance of species, while the y-axis depicts the predicted abundance by Metax. The Pearson correlation coefficient *r* and p-value are displayed. All species with either real abundance or predicted abundance above 0.1% in a sample were included (n=1419 sample-species). **c)** Abundance rank of pathogens detected by Metax detected from long-read LRI samples. If multiple pathogens are present in a sample, only the primary pathogen is counted. **d)** Predicted abundances of pathogenic and non-pathogenic taxa identified from 41 LRI samples. Only taxa with a read count over 100 are shown. The color represents the genus of each detected taxon. **e)** Taxa detected in the CAMI II pathogen detection dataset sorted by predicted abundance. All taxa with an abundance greater than 0.1% are included in the plot since Centrifuge predicted this pathogen with an abundance of 0.14%.

### Accurate detection of likely etiological agents from clinical samples of patients with infectious diseases

To assess Metax’s ability in identifying the likely etiologic agents in infections from metagenomes, we evaluated it on multiple clinical samples where the presence of a pathogen has been confirmed with techniques such as qPCR and cultivation^16,34^. In this analysis, we also considered the order of predicted abundances for the taxa as relevant, as substantial abundance is an important indicator in infectious disease diagnostics, usually assessed via colony forming units in cultivation^44,45^.

As long-read sequencing is increasingly applied in clinical metagenomics, we evaluated the performance of Metax on long read data from patients with lower respiratory tract infections^34^, comprising 34 pathogen-positive and seven pathogen-negative samples (Materials and Methods). Metax identified the likely etiologic agent for all 34 positive samples and reported no false positives for the negative samples, achieving 100% sensitivity and specificity, respectively. Furthermore, for 88% (30/34) of these samples, the pathogen taxon was ranked as most abundant, and in the remaining samples, it was ranked as second or third most abundant (Fig. 3c). Overall, the abundance of pathogen taxa is significantly higher than that of others, accounting for nearly half of the communities (52.3 ± 31.1%, Fig. 3d). Similarly, Metax also identified the Crimean-Congo hemorrhagic fever (CCHF) virus, the likely etiological agent from a clinical sample used in the CAMI II pathogen detection challenge, and ranked it as the most abundant taxon (28.16%, Fig. 3e). Of the profilers that were also assessed for the marine dataset, Centrifuge also detected the CCHF virus, however, it assigned it a very low abundance (0.14%), reducing its visibility in downstream interpretation. In comparison, the recent marker-gene based methods did not identify the virus among reported taxa, and the k-mer sketching-based technique did not report any taxa from the data.

### Reducing false positive taxon identifications via coverage statistics

As described above, Metax calculates both the observed and expected breadth of coverage for each species. The Observed-to-Expected Breadth Ratio (OEBR) is highly informative for identifying false positive taxa. For the marine benchmark dataset, we analyzed the OEBR for all species, including those with extreme OEBR, which were subsequently filtered out of Metax’s final profiling output. Binning species by OEBR revealed that those with an OEBR of around 1 are almost exclusively correct (true positives, TP), while bins with the OEBR deviating further from 1 exhibit high fractions of false positive (FP) predictions of all predictions (FDR, false discovery rate, Fig. 4a). The OEBR and the breadth of chunk coverage (Bc) also reported by Metax together are powerful metrics that complement each other; species whose observed breadth closely matches expectation (OEBR ≈ 1) and with high Bc are typically correct (low FDR, Fig. 4a), while those with large OEBR deviations can be confidently filtered out as false.

**Fig. 4.**
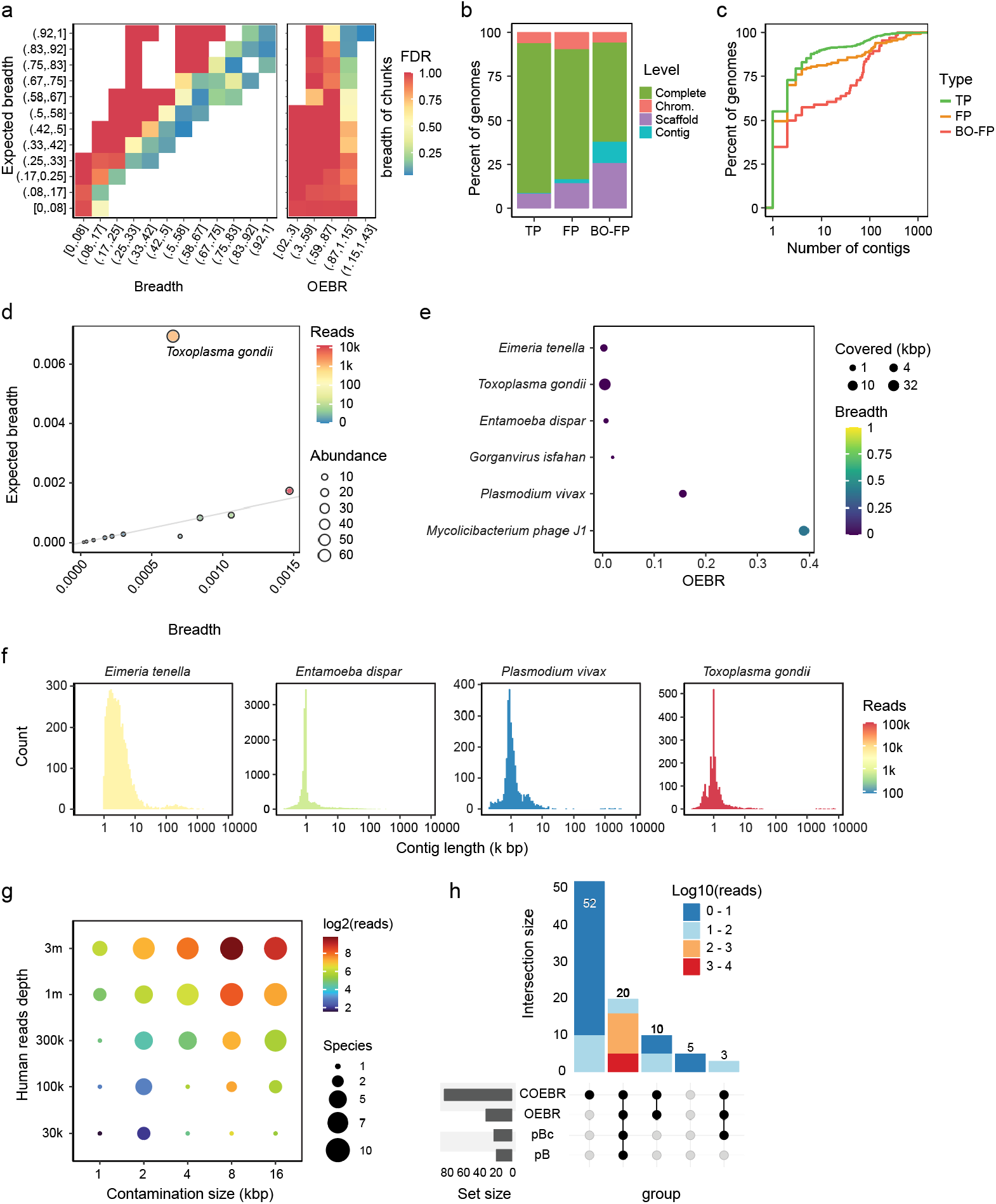
Analysis of Metax’s FP identification capability. **a)** False discovery rate across observed and expected breadth of coverage bins. The heatmap at the right, OEBR versus Bc, shares the same y-axis range and the color code with the one at the left. **b)** The genome assembly level and assembly quality of TP and FP species. BO-FP represents breadth of coverage outlier FP. These are FP species that show a significant discrepancy between their observed and expected breadth from Metax. **c**) Comparison of genome assembly fragmentation across reference genomes of TP, FP, and BO-FP (OEBR <0.75) taxa. **d)** *Toxoplasma gondii* was detected in a clinical ascites sample sequenced with Nanopore technology with low OEBR and presence likelihood by Metax. **e)** False positive microbial taxa identified by Metax by an extreme OEBR (x-axis) from the “human genome in a bottle” (GIAB) sample. The size indicates the total bases of the species’ reference genome being covered by reads. The color shows the observed breadth of coverage. **f)** The reference genome assemblies are highly fragmented for false positive taxa shown in e). The two viruses were not displayed because the recent NCBI RefSeq update no longer included their assemblies. **g**) Detection of FPs species across contamination sizes and sequencing depths for *in silico* created contaminated microbial genome assemblies. Bubble size indicates the number of species detected when human WGS reads (30 k–3 M) were mapped to bacterial genomes containing human inserts of varying lengths (1–16 kbp); color represents the log_2_ number of supporting reads. **h)** Overlap of FPs identified by different Metax metrics. The UpSet plot shows intersections among species flagged as FPs by OEBR, chunk based OEBR (COEBR), probability of presence based on breadth (pB), probability of presence based on breadth of chunks (pBc). Bar height indicates the number of FPs identified by each metric combination, and color represents the log_10_-transformed number of supporting reads. For b) and c), only taxa with supporting reads more than 1000 were considered.

To assess the impact of reference genome quality on species detection, we compared assembly quality levels (“Complete Genome” > “Chromosome” > “Scaffold” > “Contig” based on RefSeq definition) for genomes representing either TP, FP, or breadth-outlier FP (BO-FP) species with an extreme OEBR value (Fig. 4b-c). This suggested that FPs—especially those BO-FPs—are much more likely to originate from species with low-quality, fragmented genome assemblies in reference sequence collections used by profilers, and consequently, that genome assembly quality is an important factor for achieving high profiling accuracies at species level.

### Suggestion of false positives arising from contaminated reference genome assemblies

In the detection of pathogens from human clinical samples, which often have low microbial biomass, accurate distinction between human and microbial sequence data is essential. When profiling ascites samples from cirrhosis patients sequenced in-house with Nanopore long read sequencing technologies, Metax identified *Toxoplasma gondii* as a likely false positive (OEBR < 0.1, probability of presence < 10^−10^, Fig. 4d). Investigation of the read alignments revealed that all supporting reads mapped to a single short contig in the *T. gondii* RefSeq assembly. In contrast to a true *T. gondii* infection—where reads would more uniformly cover the parasite genome (OEBR ≈ 1) — the low observed breadth here suggests that rather than actually stemming from *T. gondii*, the contig comes from another source.

We hypothesized that it could originate from uncharacterized human repeat regions (e.g., telomeric or pericentromeric repeats), structurally variable loci, and polymorphic insertions, which are notoriously difficult to assemble^46^ and are only partially represented even in the most complete human genome assemblies^47,48^. If such a region were missing from human genome assemblies and included elsewhere, reads originating from there would persist in the dataset, even after reference-based host removal, and lead to false discoveries. To confirm, we applied Metax to the Genome In A Bottle (GIAB) whole-genome shotgun dataset^49^, which consists exclusively of human DNA. Indeed, using the same microbial reference database for profiling, both Metax and Centrifuge identified *T. gondii* alongside four other non-human taxa (*Entamoeba dispar, Plasmodium vivax, Eimeria tenella*, and *Gorganvirus isfahan*). Metax identified all five false positives (OEBR < 0.2; p ≪ 10^−10^) and additionally flagged *Mycolicibacterium phage J1* (OEBR < 0.5, Fig. 4e), driven by partial genome matches. Intriguingly, of these taxa, three are protozoa that reside in or infect humans and their reference genome assemblies are of very low quality and highly fragmented (Fig. 5f). This suggests that uncharacterized human sequences contaminate these reference genomes in NCBI, a phenomenon recently reported to produce artifactual microbial signals^50^. The reference genome sequences of the two phages (*Gorganvirus isfahan, Mycolicibacterium phage J1*) have been labelled “unverified source organism” in NCBI recently.

**Fig. 5:**
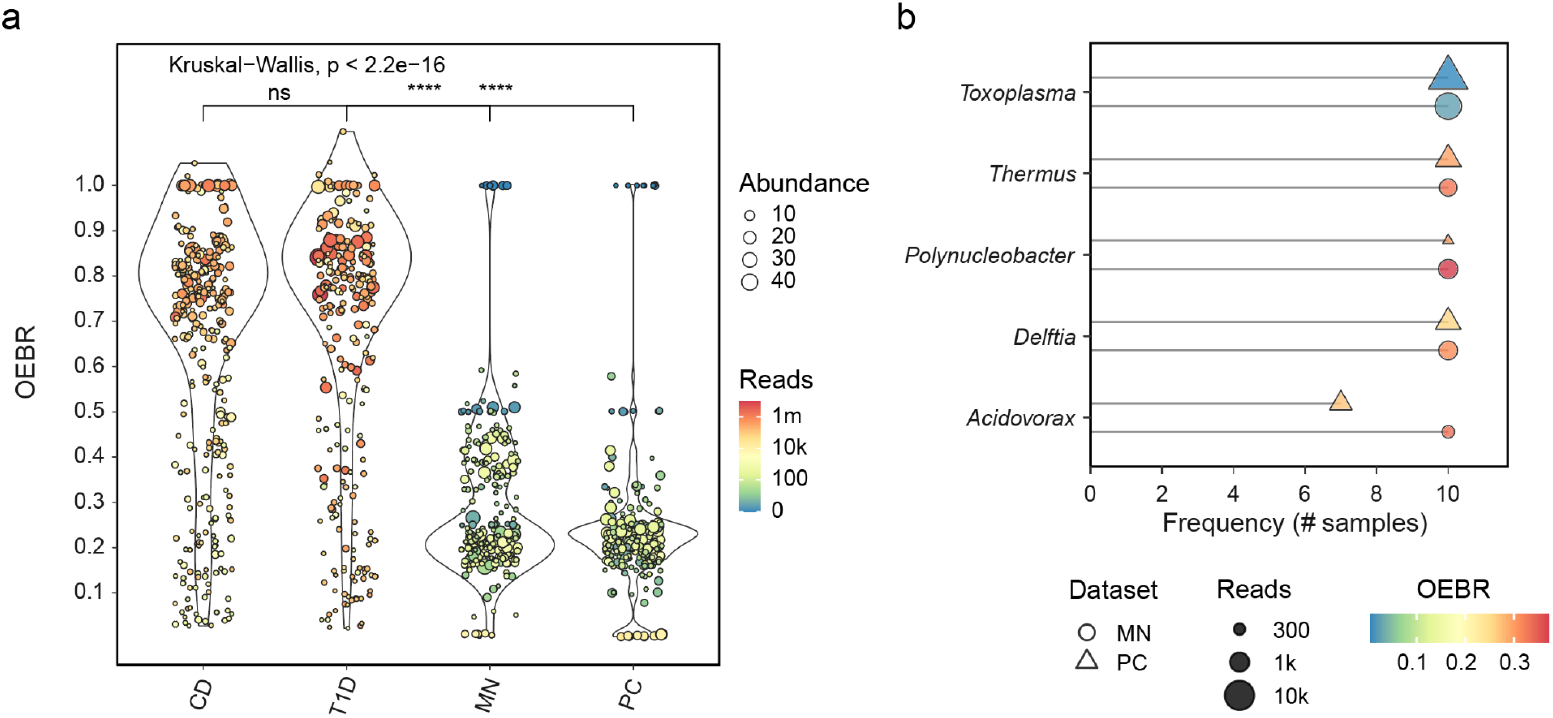
Metax identifies likely reference genome contamination, kitome from tumor microbiome datasets. **a)** OEBR for taxa predicted by Metax for four data sets without applying the probability filtering in gut and tumor microbiome datasets. **b)** The top five genera by read count “detected” in MN and PC datasets. For d. and e., only taxa with an abundance >0.5% and expected breadth >0.001 are shown in this plot for clarity. The width of the violin is weighted by abundance.

We also generated a synthetic benchmark reference set in which human DNA fragments were deliberately inserted into bacterial reference genomes. A total of 50 bacterial or archaeal genomes with genome size between 3 to 4Mb, each from a distinct species, were used; ten genomes each were spiked with human sequences of increasing size from 1 kb to 16 kb (2^n^ series). We choose the contamination size below 16k because the recent systematic study by NCBI shows that 97% of contaminant sequences in bacterial genomes are shorter than 10kbp^51^. Human whole-genome sequencing datasets at varying depths (30 k, 100 k, 300 k, 1 M, 3 M reads) were then profiled with these contaminated references using Metax. Because all reads originated from the human genome, any bacterial taxa detected were false positives induced by contamination. We observed that the larger the contaminant size and the more the human reads in the sample, the more frequently the species represented by the contaminated genome were initially detected (Fig. 4g). Metax successfully flagged 94% (85/90) of these as FPs via coverage-informed statistics (OEBR < 0.75; COEBR<0.75; p-value based on B >1e-5; p-value based on Bc >1e-5, Fig. 4h): Even as sequencing depth decreased, human contamination-derived signals could be identified, demonstrating precision under realistic and challenging conditions.

### Extreme OEBR reveals microbial signal from contaminants and kitome in tumor microbiome data

Motivated by recent debate over likely artifactual microbial signals in tumor “microbiome” studies ^52^, we leveraged Metax’s statistical framework to search for evidence for erroneous microbial signals in such datasets. Using two gut cohorts—Crohn’s disease (CD)^53^ and type 1 diabetes (T1D)^54^ as controls, we compared their OEBR distributions to those from the respective Melanoma and NSCLC-Adeno (MN) and pancreatic cancer (PC) tumor samples^55^. In the two gut microbiome datasets, taxa with OEBR >=0.75 accounted for more than 65% of the taxonomically classified reads. In both CD and T1D, the abundance-weighted densities are tightly centered around OEBR ≈ 1. Kruskal–Wallis and Wilcoxon testing confirmed no significant difference between CD and T1D, but highly significant differences (p < 2.2×10^−16^) when either gut cohort was compared to the tumor datasets (Fig. 5a). Notably, in the tumor microbiome data, the only species that was confidently identified by Metax (breadth of chunks ≥0.01 and OEBR ≥0.75) was *Aliivibrio fischeri* (in 70% of samples), which is a sequencing spike-in control. It was detected in all 20 samples; however, in three the read count was too low to support statistical confidence, and in another three the OEBR fell below 0.75. Additionally, *Thermus* species were the most frequently detected taxa with extreme OEBR values in the tumor microbiome datasets. Given that *Thermus* DNA is a well-known “kitome” contaminant introduced during library preparation^56,57^, its pervasive detection here almost certainly reflects reagent-derived DNA fragments rather than true presence. Interestingly, *T. gondii* —flagged for reference genome contamination in our above analyses—also appeared frequently with an extreme OEBR. Moreover, more species from genera that were frequently reported as originating from DNA extraction kits, PCR reagents, and laboratory environments were detected but with extremely low OEBR, including *Delftia*^57,58^, *Acidovorax*^19,57,59^, *Polynucleobacter*^19,57,58,60^ (Fig. 5b). These results demonstrate the capacity of Metax to suppress spurious microbial hits in tumor data^50^, while also detecting commonly reported contaminant taxa.

### Microbial marker discovery and phenotype prediction based on abundance profile

To demonstrate Metax’s capacity to predict the presence and abundances of taxa from multiple domains and of viruses, we analyzed an oral metagenomic dataset consisting of 317 samples from 91 subjects^61^, with 77 from healthy sampling sites, 111 from mucositis, and 126 from sites with peri-implantitis. Metax determined that viruses contribute to a significant portion of the community composition (Fig. 6a). Among the 18 communities dominated by viruses (accounting for at least 50% of the total abundance, Fig. 6b), the virus family Redondoviridae was the most abundant, followed by Papillomidaviridae. Redondoviridae were also the most prevalent, discovered in 23% (73) of the samples, followed by Papillomaviridae, which were found in 17% (54) samples (Fig. 6c). Both have a very small genome below 10k, making it challenging for profilers to detect them. A SMACOF-based multidimensional scaling (MDS) analysis of the taxon profiles clearly separated healthy sample sites from sites with peri-implantitis (Fig. 6d). Differential abundance analysis using ANCOM-BC^62^ identified 14 species significantly (adjusted *p*-value < 0.05) more abundant in peri-implantitis and another 14 were more abundant in healthy sites (Fig. 6e). Notably, all members of the red complex—well-established contributors to periodontal disease^63,64^—as well as one Redondo virus were identified as significantly more abundant in peri-implantitis.

**Fig. 6:**
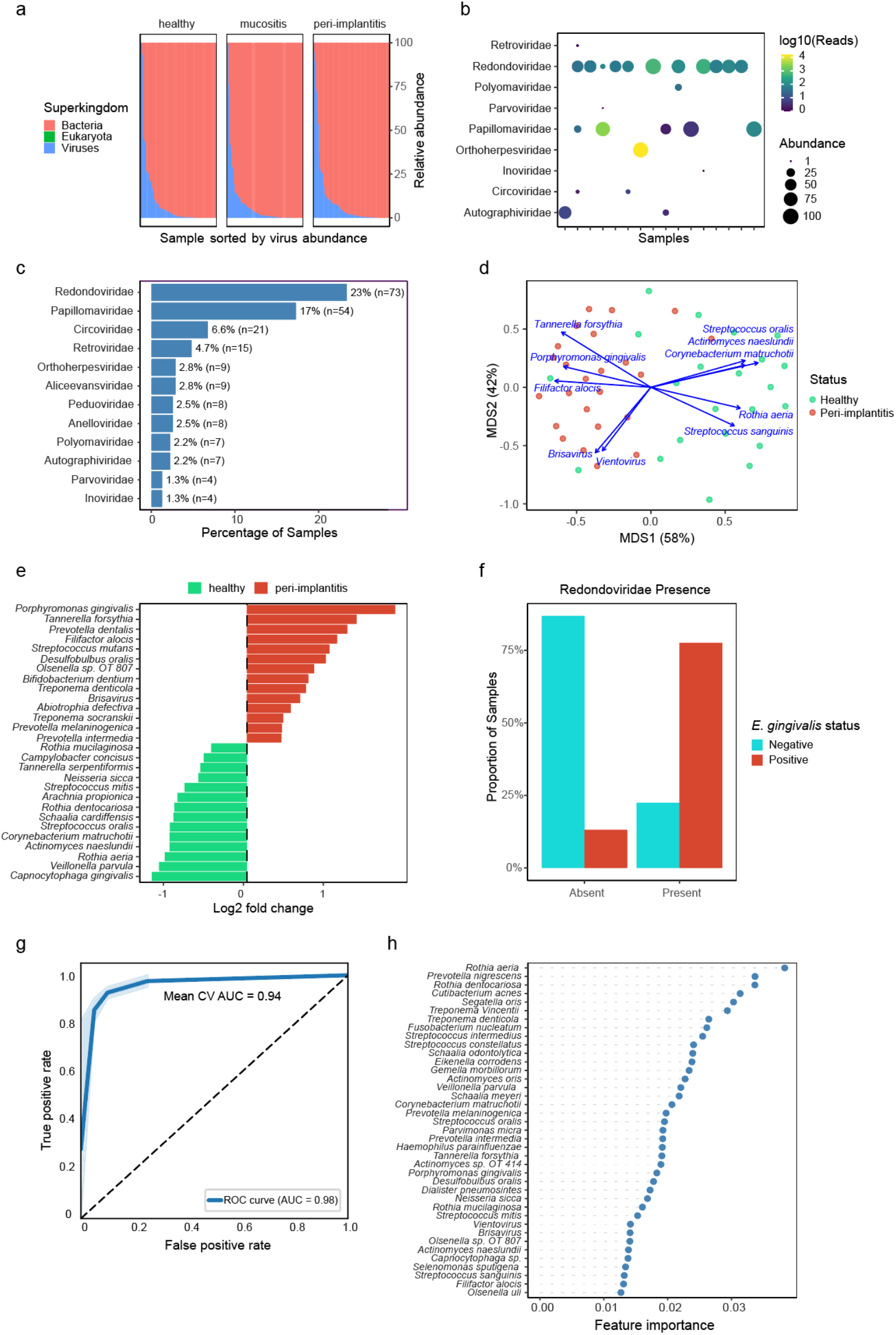
Analysis of an oral microbiome cohort with Metax. **a)** Abundance of bacteria, eukaryotes and viruses across samples. **b)** The abundance and read count of virus families in virus-dominated samples.**c)** The prevalence of different virus families in all samples. **d)** MDS visualisation of samples from healthy sites and those with peri-implantitis. **e)** The microbial species that were differentially abundant in health and peri-implantitis. **f)** The co-occurrence of *E. gingivalis* and Redondoviridae in all samples. **g)** The AUCs of the 10 cross validation folds and final classifier. **h)** The taxa that contributed to the classifier to distinguish health from peri-implantitis.

A recent study revealed that *Entamoeba gingivalis* is the host of Redondo viruses^65^. *E. gingivalis* is a parasite, which often actively resides in the periodontal pocket of the patient with oral inflammatory conditions^64,66^. However there is no *E. gingivalis* genome available for profiling, only its 18 srRNA is known. Therefore we mapped reads of all samples against the 18 srRNA gene of *E. gingivalis*. Comparing the presence of the *E. gingivalis* 18 srRNA gene and the abundance of Redondo virus predicted by Metax, we observed a strong, significant association between these two (OR = 22.3; 95% CI 9.5–56.6; p < 2.2 × 10^−16^, Fisher’s exact test; Fig. 6f). Moreover, in eight samples, Metax detected both TM7 species and their putative host bacteria *Actinomyces*. TM7 bacteria, members of the Candidate Phylum Radiation, possess highly reduced genomes—lacking many core metabolic and biosynthetic pathways— and thus depend on host bacteria to supply essential functions^67,68^. Metax detected both TM7 and its host, thus facilitating investigation of their obligate symbiosis and in general of microbial interactions across or within domains.

A Random Forest model based on the Metax taxon profiles distinguished the peri-implantitis sites from healthy sites with an average Area under the Curve (AUC) of 0.94 in 10-fold cross-validation and a best-model AUC of 0.98 (Fig. 6g). Strikingly, among the most important taxa for the classifier decision, all red complex members, most orange complex members (e.g., *Prevotella intermedia, Fusobacterium nucleatum*) as well as Redondo virus were included (Fig. 6h). These findings illustrate that Metax’s high-fidelity, cross-domain taxonomic profiles can yield biologically coherent disease-associated signatures.

## Discussion

Taxonomic profiling is a common task in metagenomics and basis for insights into the structures of microbial communities. In attaining high quality profiles, key challenges are the profiling low ranking taxa, as well as accurate assignment beyond bacterial taxa, such as for viruses and archaea^16^. From a technical perspective, intrinsic mapping ambiguity driven by short reads and shared genomic regions among related taxa^18^, fragmented or contaminated reference assemblies^21,51,69–71^, and reagent-derived DNA contamination^60^ can cause issues. These issues are further amplified in low-biomass or host-dominated samples, where microbial reads are sparse and background noise can overwhelm true biological signals^23^. Here, we describe Metax, a taxonomic profiler combining genome-read coverage statistics from mapping metagenome data to reference genome collections with a probabilistic EM framework, which efficiently addresses these challenges. Metax supports both short- and long-read data with high computational efficiency. While it applies the NCBI taxonomy, it can be easily reconfigured to alternative frameworks—such as GTDB^72^ for prokaryotes or ICTV^73^ for viruses—using user-provided taxonomy dump files.

In extensive evaluations across complex, deeply sequenced gut microbiomes, Metax consistently delivered accurate taxon profiling and abundance estimation across sequencing depths, and taxonomic domains, including viruses. Performance benefits with key metrics relative to techniques based on k-mer sketching, marker genes or k-mer matching are particularly evident at the species-level. Furthermore, substantial benefits for viral taxon detection were seen, which is particularly notable, as these groups are typically challenging due to small genome sizes, high divergence^74–76^. Metax also maintained runtime efficiency comparable to computationally lightweight approaches^15,28^, completing profiling of each 2 Gb sample within minutes.

For low-biomass clinical samples, where shallow sequencing depth and/or high host background typically result in substantial reductions in taxonomic profiling accuracy^19,23,57,77^. Metax retained robust taxon detection at depths as low as 1,000 reads per sample, and the lowest abundance estimation errors across all taxa groups. Metax was the only profiler identifying the etiologic pathogen in the CAMI II pathogen challenge data and achieved 100% sensitivity and specificity on Nanopore long-read clinical samples from patients with lower respiratory infections, and ranking the pathogen as most abundant in nearly all positive cases. These results highlight Metax’s suitability for translational and diagnostic applications, where high specificity and robustness to noise are essential^4,23^.

We found that coverage-based metrics provided by Metax effectively separate true microbial signals from artifacts arising from reference contamination, conserved genomic regions, or mobile genetic elements. Taxa with extreme discrepancies between observed and expected breadth were almost exclusively false positives. When profiling human whole-genome sequencing data—where no microbial DNA is expected—several microbial reference genomes produced false protozoan hits, likely due to uncharacterized human sequences embedded in their assemblies. Metax correctly flagged all of these as false positives, demonstrating its ability to recognize and filter out artifacts caused by contaminated or misassembled reference genomes. Consistent with this observation, our simulation experiment, which introduced defined human DNA fragments into bacterial reference genomes, demonstrated that Metax consistently identified the resulting contamination-driven false positives, even at low sequencing depth. As public reference repositories continue to grow faster than curation efforts, such contamination-aware detection becomes increasingly essential for reliable metagenomic profiling.

Application to tumor “microbiome” datasets—where microbial presence remains highly debated^52^—further demonstrated Metax’s ability to suppress artifactual signals. In gut microbiomes from Crohn’s disease and type 1 diabetes cohorts, the obtained genome coverage signals of taxa were consistent with genuine community structure. In contrast, coverage distributions for taxa detected in plasma samples from MN and PC patients were indicative of noise. The only species confidently retained across tumor samples was *Aliivibrio fischeri*, a deliberate spike-in control. These results align with recent evidence that most microbial signals in tumor sequencing derive from reagent contamination and misassembled reference genome rather than true intratumoral microbiota^50^, and illustrate Metax’s ability to prevent erroneous biological interpretation in such contentious low-biomass contexts.

Finally, analysis of an oral metagenomic cohort illustrated how biological insights can be obtained by analysing Metax’s high-fidelity, cross-domain profiles. Community structure clearly separated healthy and peri-implantitis sites, and differential abundance analysis recovered well-established oral disease signatures, including the red and orange complex bacteria^63,64^. Notably, Metax also identified substantial viral contributions to these signatures, including Redondoviridae and Papillomaviridae, which are difficult to detect due to their extremely small genomes. Additionally, the cross-domain profiles delivered by Metax capture both the recently described Redondoviridae–*E. gingivalis* interaction^65^ and the established TM7–Actinomyces association^67,68^, illustrating its ability to recover ecologically meaningful relationships involving ultra-small or obligately symbiotic taxa.

Together, our findings show that Metax offers a robust, accurate solution for cross-domain taxonomic profiling and contamination removal across a wide spectrum of sample types and sequencing depths. Unifying coverage-informed presence probability estimation with EM-based abundance refinement overcomes issues caused by ambiguous read mapping, reference genome misassemblies, and reagent contamination in taxonomic profiling. These properties make Metax a powerful technique for microbiome research and clinical metagenomics, and for improving genome assembly qualities. Future directions of where this approach could excel include strain-resolved models, profiling of RNA viruses, and detecting contaminated or misassembled reference genomes at database scale.

## Material and Methods

### Sequence Alignment and parameter settings

Metax employs the Modular aligner^24^ to map reads against a taxonomically annotated reference genome database. The reference genome sequences include the taxonomy ID in their header lines. For each alignment, Metax computes the gap-compressed sequence identity between the read (query sequence) and the reference, as well as the mapped read length and fraction based on the CIGAR string. An alignment with identity, mapped length, and mapped fraction above defined thresholds is considered a valid alignment. For short-read data, the default identity, mapped length and fraction cutoffs are set to 0.97, 50, 0.8, respectively. On short reads data for pathogen detection, an identity threshold of 0.95 and a mapped fraction of 0.7 were used. For long-read data, the defaults are 0.86, 250, 0.6, respectively. Species with OEBR <0.75 or >1.5 and p-value <1e-5 were filtered out in the final taxonomy profile. To robustly assess read-mapping uniformity—particularly for taxa with sparse read counts—we introduced a breadth-of-chunk coverage metric, which divides each reference genome into equal-length segments (“chunks”) and computes the fraction of those segments covered by at least one read. For long-read datasets, genomes are partitioned into approximately 20 chunks, whereas for short-read datasets, genomes are split into 1,000 chunks to capture finer-scale coverage patterns. Metax offers a host-filtering option: when a host taxon is specified, non-host-associated viruses are omitted, yielding a streamlined taxonomic profile that highlights the true pathogens.

### Calculation of abundance rank error

Existing abundance evaluation metrics such as Bray–Curtis dissimilarity, Pearson correlation, and Weighted UniFrac quantify overall abundance or abundance weighted phylogenetic similarity, but do not assess whether taxa are ranked in the correct order of abundance. The Bray–Curtis distance can be low even if the dominant taxa are misplaced, which is misleading in pathogen detection and other clinical metagenomic analyses, where the etiological agents are identified from prevalent taxa in infected samples. For these applications, the order of abundance is more biologically meaningful than the precise abundance values themselves. In such cases, even moderate abundance deviations are less critical than errors in taxon ranking, especially when the top-ranked organism determines diagnostic interpretation or treatment decisions. To quantify the disagreement in the ordering of taxa between the predicted and ground-truth profiles, we defined the abundance rank error (ARE).

For each sample and taxonomic rank, taxa are sorted by decreasing abundance in both profiles. Let 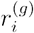 and 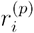 denote the rank positions (1 = most abundant) of taxon *i* in the ground-truth and predicted profiles, respectively, and let *n*_*g*_ and *n*_*p*_ be the numbers of taxa in each.

Each taxon in the ground truth is assigned a rank-based importance weight

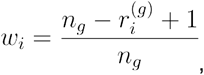

so that higher-ranking taxa contribute more strongly to the total error.

The total rank error E is computed as the sum of three components:

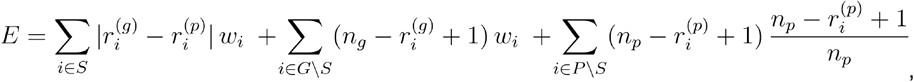

where S is the set of shared taxa, G and P are taxa only in the ground-truth and predicted profiles, respectively.

The first term penalizes rank mismatches for shared taxa, the second penalizes taxa missed by the profiler, and the third penalizes taxa predicted but absent in the ground truth.

The result is normalized by the theoretical maximum error (no taxa overlap):

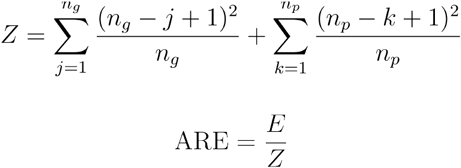

The ARE ranges from 0 (perfect rank concordance) to 1 (maximal disagreement). Smaller values indicate that dominant and subdominant taxa are ranked similarly between profiles, whereas large values arise when major taxa are missing, falsely predicted, or misordered. By focusing on rank consistency, the ARE complements conventional abundance- and phylogeny-based distances, providing a biologically meaningful measure of profiling accuracy—especially relevant for pathogen detection, low-biomass samples, and other scenarios where the identity of the top-ranking taxa carries the most diagnostic importance.

### Benchmark datasets

#### Simulated multi-domain gut metagenomic dataset

To evaluate cross-domain profiling accuracy, we simulated a human gut metagenomic dataset containing bacterial, archaeal, fungal, and viral taxa designed to resemble the compositional and taxonomic structure of real gut microbial communities. Ten representative gut microbial communities were derived from previously published metagenomic abundance profiles^32^ to serve as the prokaryotes backbone for simulation. Additional microbial domains were incorporated to reflect realistic multi-domain diversity: viruses (∼49% of total abundance), fungi (∼1%), composed of 80 bacteriophages infecting the most prevalent bacterial hosts in these communities and 20 human viruses frequently detected in the gastrointestinal tract. Virus–host associations were curated from the Virus-Host DB^78^ to ensure biological plausibility. A trace amount of human DNA (0.002%) was also included to emulate host contamination typically observed in gut metagenomic samples. Relative abundances across domains were normalized to predefined proportions (bacteria + archaea = 50%, viruses ∼ 49%, fungi ∼ 1%, human ∼ 0.002%) following a log-normal/Dirichlet hybrid distribution to reproduce natural long-tailed abundance patterns. Shotgun sequencing reads were simulated using CAMISIM^79^ with the Illumina HiSeq error model, producing approximately 2 Gbp of paired-end reads per sample (2 × 150 bp).

#### Clinical datasets simulation with different sequencing depth

To evaluate profiler performance under different biomass conditions, we in silico generated metagenomic datasets modeling ascites and cerebrospinal fluid (CSF) communities at four distinct sequencing depths. First, ten ascites and ten CSF communities were designed by selecting taxa commonly reported in clinical studies: each ascites community comprised three bacterial species, one fungal species, and two viral species, while each CSF community comprised three bacterial species, two fungal species, and two viral species^39^. The abundance of each species in the communities was randomly generated using an in-house script while limiting the maximum abundance of fungi due to their larger genome size. For each community, we produced four replicate samples at depths of 1,000 (1 k), 10,000 (10 k), 100,000 (100 k), and 1,000,000 (1 M) microbial reads to span the range from ultra-shallow to moderate depth. Reference genomes for these species used for simulating reads were obtained from NCBI RefSeq. Sequencing reads were simulated using ART (v2.5.8)^80^ with the Illumina HiSeq 2500 error model, generating 150 bp paired-end reads. The same reference genomes set was used in Metax and other profilers except for marker-gene methods (MetaPhlAn4, mOTUs3) which employed their own curated marker sets. Phages were excluded from all profiler outputs.

#### CAMI II marine dataset

We evaluated the performance of Metax on short and long reads data from this CAMI II dataset and compared it with other profilers including, Bracken, CCmetagen^83^, Metalign, MetaPhlAn2, mOTUs2, MetaMaps^43^ and Centrifuge^31^ and latest MetaPhlAn4 and mOTUs4. MetaMaps and Centrifuge were only evaluated on long reads data. Since Metax supports both short and long reads (Nanopore, PacBio) data, we evaluated it on both short (noted as Metax_short) and long (Metax_long) reads data from CAMI II marine dataset. KMCP was not included due to its long runtime and high computational demands^84^.

#### Pathogen detection datasets

CAMI II pathogen detection dataset and the LRI dataset were used to evaluate Metax’s accuracy for pathogen detection on both short and long-read data. The CAMI II pathogen detection dataset was derived from a patient suffering from Crimean-Congo Hemorrhagic Fever (CCHF). A blood sample was taken from the patient, and the extracted RNA was sequenced using Illumina technology. The LRI dataset comprises 41 lower-respiratory-tract metagenomic samples sequenced with long-read Oxford Nanopore technologies^34^. Out of these, 34 samples were pathogen positive as confirmed through culture and/or PCR and seven are negative. In the original study, 13 bacterial species were designated as LRI pathogens^34^. A relative abundance cutoff 5% was applied to eliminate background noise in the data, as in the originating article. For both datasets, non-human associated viruses were removed from the taxonomy profile of all profilers.

### Evaluation of reference contamination and false-positive detection

To evaluate Metax’s ability to identify false positives arising from contaminated reference genomes, we constructed a controlled benchmark dataset simulating contamination between human and prokaryotes sequences. A total of 50 complete bacterial or archaeal genomes with genome size between 3-4Mbp, each representing a distinct species, were randomly selected from the RefSeq database. To emulate reference contamination, we inserted fragments of human genomic DNA of increasing length into these bacterial genomes. Specifically, human sequences with lengths of 1 kb, 2 kb, 4 kb, 8 kb, 16 kb were extracted from different non-overlapping regions of the T2T human genome assembly. Ten genomes were assigned to each contamination length, resulting in a reference collection that incorporated human fragments of defined sizes. The modified bacterial references were concatenated into a single FASTA file as reference for profiling. To generate the test data, we used the GIAB data and subsampled it to five sequencing depths—30 k, 100 k, 300 k, 1 M, and 3 M reads—to assess sensitivity under varying host background levels. Each read set was then profiled against the contaminated reference collection using Metax. Metax was evaluated for its ability to detect and flag these false positives using its coverage-informed metrics, including the OEBR, chunk based OEBR (COEBR), probability of presence based on Breadth (pB), and probability of presence based on Breadth of chunks (pBc).

### Comparison of gut microbiomes and cancer microbiomes

We obtained metagenomic sequencing data from four published studies: gut microbiome samples from Crohn’s disease patients^53^ and the Type 1 Diabetes Diabimmune study^54^, as well as blood plasma–derived microbiome data from melanoma and NSCLC-Adeno and prostate cancer patients^55^. For Fig. 5a, to improve clarity and ensure direct comparability, we visualized the results from the 10 samples with the highest read counts from each dataset. The Kruskal–Wallis test and Wilcoxon test were used to assess whether OEBR distributions differed significantly among the datasets.

### Oral microbiome cohort analysis

Short read metagenomic data for 317 subgingival-plaque samples (77 healthy, 111 mucositis, 129 peri-implantitis) were obtained from a published oral microbiome study^61^. Metax-derived species-level relative abundances were transformed into a sample × taxon matrix, which was then subjected to two-dimensional ordination using the Scaling by MAjorizing a COmplicated Function (SMACOF) algorithm implemented in R library smacof^85,86^ on Euclidean distances. Differential abundance between healthy and peri-implantitis sites was assessed with ANCOM-BC (v2.8.1)^62^, applying a Benjamini–Hochberg–corrected false discovery rate (FDR) cutoff of 0.05 to identify significantly altered taxa. To detect *E. gingivalis*, which lacks a complete genome in our reference set, reads were aligned to its 18S rRNA gene (GenBank: D28490.1) using BWA-mem2 (v2.2.1)^87^. Co-occurrence between Redondoviridae and *E. gingivalis* was evaluated by Fisher’s exact test on presence/absence (presence defined as ≥2 mapped 18S reads). Finally, a random-forest classifier was trained on Metax species abundances (scikit-learn v1.7.1 with default parameters) using ten-fold cross-validation; model performance was summarized by average area under the ROC curve (AUC), and feature importance was ranked by mean decrease in Gini impurity.

### Reference databases

For the benchmarks on the CAMI II datasets (marine, and pathogen detection), we utilized the genomes available from the NCBI RefSeq database as provided on the CAMI II website (snapshot of 8 January 2019). This reference genome set comprises 18,098 bacterial, archaeal, and viral genomes and was used for all profilers except MetaMaps and the marker-gene–based methods. Due to compatibility issues when building a MetaMaps database from this collection, we instead employed the default database provided in the MetaMaps GitHub repository. For all other analyses, we used the genomes from the snapshot of the RefSeq database captured on August 10, 2022. In total, 33,143 genomes from bacteria, archaea, viruses, fungi, protozoa and homo sapiens are included in the reference database. Metax_newdb in the gut metagenomic dataset evaluation used 33,533 genomes from an updated RefSeq database from July 30, 2025. MetaPhlAn4 and mOTUs3 were run with their own up-to-date marker-gene databases.

### Hardware and software for benchmarks

The performance metrics were calculated with OPAL^81^. Snakemake^82^ was used to record the runtime and memory consumption on a server equipped with 40 CPU cores (Intel Xeon Processor (2.6 GHz)) and 256GB of memory.

## Software and data availability

Metax is available from https://github.com/hzi-bifo/Metax, and can be installed with Conda. The ground truth and profiler generated taxonomy profiles for this study can be accessed with DOI: 10.5281/zenodo.17776358.

## Acknowledgements

We thank Gary Robertson for IT support and Dr. Mohammad-Hadi Foroughmand-Araabi for advice on statistical formulations. Z.-L. Deng was partly funded by the Deutsche Forschungsgemeinschaft (DFG, German Research Foundation) under Germany’s Excellence Strategy - EXC 2155 - project number 390874280 and the German Center for Infection Research (DZIF, Grant TI BBD 12.002). N. Safaei was partly funded by zukunft.niedersachsen, the joint science funding program of the Lower Saxony Ministry of Science and Culture and the Volkswagen Foundation.

