## Supplementary Fig. 1 for "Metax: A Coverage-Informed Probabilistic Framework for Accurate Cross-Domain Taxon Profiling"

### Supplementary data

#### Figures

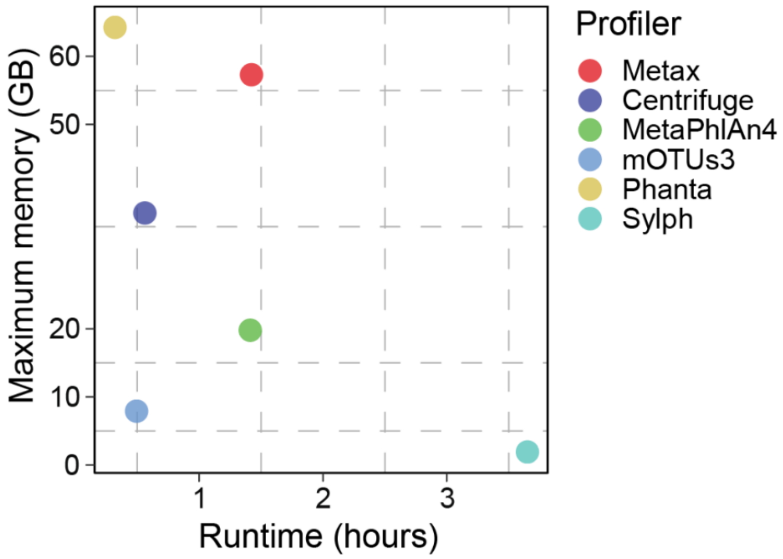


**Fig. 1: The total runtime and maximum memory consumption of profilers on simulated gut metagenome dataset.** The runtime indicates the total hours required to complete the entire dataset which is recorded by Snakemake.
